# Characterisation of aphid antixenosis in aphid-resistant ancestor wheat, *Triticum monococcum*

**DOI:** 10.1101/2024.07.01.601505

**Authors:** Alexander N. Borg, József Vuts, John C. Caulfield, David M. Withall, M. John Foulkes, Michael A Birkett

## Abstract

**BACKGROUND:** Due to the increasing presence of insecticide resistance across cereal aphid populations, new aphid management strategies, including the engineering of host resistance to aphids into commercial wheat varieties, are required. Previous studies have identified ancestor wheat, *Triticum monococcum* accessions MDR045 and MDR049, with resistance against the grain aphid, *Sitobion avenae*. To test the hypothesis that resistance can be accounted for by antixenosis (reduced attractiveness of host plants) *via* the release of repellent volatile organic compounds (VOCs), we explored the response of *S. avenae* to MDR045 and MDR049 following *S. avenae* herbivory, using behaviour and electrophysiology experiments.

**RESULTS:** In four-arm olfactometry assays, alate *S. avenae* showed aphid-density dependent reduced preference to VOC extracts from *T. monococcum* MDR045 and MDR049. By contrast, alate *S. avenae* showed aphid-density dependent increased preference to extracts from hexaploid wheat, *T. aestivum* var Solstice and *T. monococcum* MDR037. Coupled gas chromatography-electroantennography (GC-EAG), using the antennae of alate *S. avenae*, located 24 electrophysiologically active compounds across all tested accessions. Synthetic blends created from 21 identified EAG-active compounds confirmed bioactivity of corresponding VOC extracts in four-arm olfactometry assays against alate *S. avenae*.

**CONCLUSION:** Our data suggest that resistance of *T. monococcum* MDR045 and MDR049 to *S. avenae* can be at least partially accounted for by antixenosis, through antennal perception of specific repellent VOC blends induced by *S. avenae* feeding behaviour.

## 1 Introduction

Wheat, *Triticum aestivum* L. (Poaceae), is a major global food crop, which has consistently been in the top five most-produced commodities worldwide between 2015-2021, forming part of the staple diet of a large portion of the global population.^1^ Within the UK, wheat had the highest yield production compared to all other crops in 2021, of 14.5 million tonnes valued at £2.7 billion, showcasing its importance as a food crop within the UK.^1^ Aphids (Hemiptera: Aphididae) are major agricultural pests on wheat and other cereals, causing significant damage by reducing nutrient and assimilate availability via phloem feeding, by viral transmission, and by reducing photosynthesis due to aphid honeydew enabling saprophytic fungal growth on leaves.^2,3^ These combined factors, particularly the viruses that aphids transmit, make aphids the most economically damaging cereal pests, leading to yield losses of 5-80% in wheat.^4^

Aphid infestations have mainly been controlled through the use of insecticides.^5^ However, the increasing presence of insecticide resistance across aphid populations, the initial and re-registration costs of insecticides and the banning of working insecticides due to their environmental impact is reducing the options farmers have to control these pests.^5–7^ This has led to new initiatives to tackle aphid infestations, one of them being investigations into naturally occurring aphid resistance across cereal lines, which can potentially be engineered into commercial wheat varieties.^8^ This approach has been successful against the Russian wheat aphid *Diuraphis noxia* Kurdjumov, with several resistance genes identified and integrated into modern wheat.^9^ However, suitable resistance traits remain lacking for the two major aphid cereal pests in the UK and northern Europe, the English grain aphid, *Sitobion avenae* F. and the bird-cherry-oat aphid, *Rhopalosiphum padi* L., both of which are vectors for barley yellow dwarf virus (BYDV)^10^.

Although aphid resistance has been identified against both *S. avenae* and *R. padi* in *T. aestivum* varieties,^11,12^ ancient wheat varieties have been found to be more resistant.^13–15^ This was observed in settling assays, where fewer *R. padi* settled on 10 of 12 tested *T. boeticum* Boiss. and *T. monococcum* L. varieties compared to *T. aestivum*, indicating that volatile organic compounds (VOCs) play a role in reduced aphid settlement.^16^ Further studies identified two promising *T. monococcum* varieties, MDR045 and MDR049, that show signs of aphid resistance against both *S. avenae* and *R. padi*.^17–20^ *T. monococcum* is a diploid ancestor wheat closely related to the A genome donor of hexaploid wheat, *Triticum urartu* Thumanjan ex. Gandilyan.^21^ In field studies, both MDR045 and MDR049 had lower aphid numbers and MDR049 had more aphid predators than *T. aestivum*.^18^ Furthermore, VOC extracts from *R. padi*-infested and un-infested MDR049 induced reduced aphid preference in four-arm olfactometry assays, with six compounds identified as potential repellent VOCs.^22^ These studies suggest that antixenosis (reduced attractiveness) plays a part in MDR045 and MDR049 resistance against *S. avenae* and *R. padi*.

The use of VOCs in integrated pest management (IPM) strategies is a well-established practice. They can be synthetic or natural and derived from either insects, such as pheromones, or plants, and are used in IPM strategies to catch or disorientate pests, recruit natural enemies or prime neighbouring plants for enhanced defence responses.^23–25^ Furthermore, VOCs, presented either *in planta* or as synthetic blends, can be incorporated into push-pull IPM strategies, reducing the need for insecticides.^25,26^ Alternatively, identification of the genetic mechanisms involved in plant VOC-mediated aphid resistance facilitates their incorporation into modern crop varieties via genetic engineering or classical breeding approaches.^4,10,27^ Genetic approaches reduce the need for pest protection products and allow for the development of a crop variety with multiple aphid resistance mechanisms, reducing the chances of the pest overcoming this resistance. This approach has been trialled with limited success through the incorporation of synthetic genes into wheat, rice (*Oryza sativa* L.) and cotton (*Gossypium hirsutum* L.). However the approach still holds promise for the incorporation of genes from more closely related species.^28–30^

Following the identification of aphid resistance in *T. monococcum* MDR045 and MDR049, we aimed to characterise the VOC-mediated aphid resistance mechanisms within these two germplasms.

VOC extracts from *S. avenae*-infested MDR045 and MDR049 were screened, alongside VOCs from aphid-infested MDR037 and *T. aestivum* Solstice (aphid-susceptible), in behavioural (four-arm olfactometer) assays with alate *S. avenae*, to assess whether bioactivity is aphid-density dependent. Coupled gas chromatography-electrophysiology (GC-EAG) and coupled GC-mass spectrometry (GC-MS) were used to locate and identify bioactive VOCs within extracts.

## 2 Materials and Methods

### 2.1 Insect and plant material

i. *S. avenae* originating from the Rothamsted farm (Hertfordshire, UK), co-ordinates: 51°48’36.8”N 0°22’34.4”W, were reared on wheat, *T. aestivum* Solstice (Advanta Seeds UK Ltd), in ventilated Perspex cages kept at 20°C, 60-70% humidity and at a 16:8h light: dark regime. Plants were replaced every two weeks to maintain an apterous aphid culture. Alates were developed when needed by allowing the culture to become overcrowded. Aphids were retrieved from cultures using a fine-haired brush and placed in clip cages when required for use.
ii. *T. monococcum* MDR037, MDR045 and MDR049 seeds were provided by the Wheat Genetic Improvement Network (WGIN) and *T. aestivum* Solstice seeds were provided by Rothamsted Research; seeds were stored at 4°C until use. Seeds were sown in Rothamsted Prescription Mix (Petersfield Products, Leicestershire, UK) and grown in controlled environment rooms at 21°C and 16:8h light: dark regime until required for use. All experiments were run using 14-day-old plants.

### 2.2 Dynamic headspace collection (air entrainment)

Fourteen-day-old plants were subjected to 0, 1, 5, 10 and 25 aphid density treatments, using apterous *S. avenae* confined on the flag leaf of each plant in clip cages.^31^ Dynamic headspace collection was carried out using air entrainment kits (Pye volatile collection kit, Kings Walden, Hertfordshire, UK) after placement of clip cages on plants, with plants enclosed in transparent cooking bags (Sainsbury’s Supermarkets Ltd, UK) secured by wire ties. Porapak Q adsorbent tubes consisting of a 4 mm internal diam. borosilicate tube filled with 50 mg Porapak Q (Supelco, Bellefonte, PA, USA) sandwiched between two glass wool plugs was used for the collection of headspace extracts. Charcoal-filtered air was passed into the cooking bag at 500 mL min^-1^ and pulled out, through the adsorbent tube, at 400 mL min^-1^, collecting VOCs emitted by the plants on the adsorbent tubing over 24 hrs. Trapped VOCs were recovered from the Porapak Q tubes by eluting with redistilled diethyl ether (750 µL, Fisher Scientific, Loughborough, Leicestershire, UK). Headspace extracts were concentrated to 100 µL under a gentle flow of nitrogen and stored at -20°C until required for use. Four replicates were carried out for each treatment using an alpha design with additional Latinization by blocks such that each treatment type would be present in every possible position of the experiment design across replicates. Following dynamic headspace collection, all plant shoots were collected, dried in an oven at 80°C overnight and weighed.

### 2.3 Aphid behaviour assays

The bioactivity of VOC extracts against alate *S. avenae* was assessed using four-arm olfactometer assays.^32^ The olfactometer consisted of 3 layers of 6 mm thick Perspex with an internal diameter of 115 mm, in which the middle layer was fashioned to consist of 4 side arms/areas at 90° angles to each other, and a central area. Each side arm narrowed at the perimeter leading to a 3 mm hole, in which glass syringes can be placed to test headspace extracts. The lower layer was lined with a 110 mm diameter Whatman® type 1 filter paper (Maidstone, Kent, UK) providing traction for the walking aphid. The top Perspex layer contained a central hole (3 mm diameter), which was connected to a vacuum pump pulling air through each olfactometer arm at a rate of 75 mL min^-1^. Prior to use, glassware was washed with Teepol, rinsed with water and acetone and baked at 130°C overnight, whilst the Perspex was washed with Teepol, rinsed with water and 70% ethanol and allowed to air-dry. The olfactometer was placed in a box lined with black paper to reduce the effect of external stimuli and illuminated from above by diffuse uniform light from two 18 W/35 white, fluorescent light bulbs. Headspace extract samples from replicates 1 and 4 from each treatment (containing the most similar GC profiles across replicates) were combined and evaporated down to 100 µL under flow of nitrogen. Ten µL of the pooled sampled, equivalent to the VOCs released by 5 plants over 20 mins, was placed on a strip of Whatman® type 1 filter paper and placed in the treatment glass syringe attached to one arm of the olfactometer. Ten µL of redistilled diethyl ether was used as a control in each of the remaining three arms of the olfactometer. A single alate *S. avenae* was placed in the central chamber of the olfactometer and left to acclimatise for 2 mins, after which the experiment was run for 16 mins, rotating the olfactometer by 90° every 4 mins. The time the aphid spent in each olfactometer arm was measured using the OLFA software (Udine, Italy, 1995). Each experiment was run at 22°C and repeated 10 times for each treatment, replacing the aphid after each run. To assess repellent activity, the olfactometer assay method was adjusted to have three treatments arms and one control arm.

Synthetic blends of identified EAG-active compounds were tested in four-arm olfactometry assays to confirm whether they are responsible for the aphid behavioural responses induced by VOC extracts. A synthetic blend for each VOC extract that produced a significant behavioural response was made in a way that the applied dose in 10 µL solution mimicked that of the corresponding VOC extract. The concentration of VOC extracts and volumes used to create synthetic blends from authentic standards are shown in Table S1 and S2, respectively. Diethyl ether (10 µL) served as control as described above.

The following experiments were conducted, in which n = aphid density:

i. *T. monococcum* MDR049 VOC extract (*S. avenae* n = 0, 1, 5, 10, 25) .v. solvent control.
ii. *T. monococcum* MDR045 VOC extract (*S. avenae* n = 0, 1, 5, 10, 25) .v. solvent control.
iii. *T. monococcum* MDR037 VOC extract (*S. avenae* n = 0, 1, 5, 10, 25) .v. solvent control.
iv. *T. aestivum* Solstice VOC extract (*S. avenae* n = 0, 1, 5, 10, 25) .v. solvent control.
v. *T. monococcum* MDR049 synthetic blend (*S. avenae* n = 1, 5, 10, 25) .v. solvent control.
vi. *T. monococcum* MDR045 synthetic blend (*S. avenae* n = 25) .v. solvent control.
vii. *T. monococcum* MDR037 synthetic blend (*S. avenae* n = 10) .v. solvent control.
viii. *T. aestivum* Solstice synthetic blend (*S. avenae* n = 10, 25) .v. solvent control.
ix. *T. monococcum* MDR049 VOC extract (*S. avenae* n = 25) .v. solvent control, repellent assay.

### 2.4 Coupled gas chromatography – electroantennography (GC-EAG) analysis

To locate compounds within behaviourally active extracts collected from the two extreme aphid density treatments (*S. avenae* n = 0 and 25), coupled GC-EAG analysis was performed using the antennae of alate *S. avenae* antennae. The set up has been previously described.^33^ Alate *S. avenae* antennae were excised at the base segment of the antenna, followed by removing the extreme tip, and suspended between two glass electrodes filled with ringer solution (without glucose). The glass electrodes were attached to Ag-AgCl wires in a way that the antennal tip was brought into contact with the recording electrode. Headspace extract (2 µL) was injected into an Agilent 6890A GC fitted with a non-polar HP1 column (50 m length × 0.32 mm inner diameter × 0.52 µm film thickness, J&W Scientific), using helium as the carrier gas and a 60 min run time starting at 30°C for 2 mins, followed by a rise of 5°C min^-1^ until 250°C. Signals from the aphid antenna were amplified (UN-06, Ockenfels Syntech GmbH, Kirchzarten, Germany) and monitored simultaneously with the GC-FID outputs using Syntech GC/EAD for Windows software (v2.3 09/1997). GC peaks were deemed to be EAG active if a response was elicited in at least 3 replicate runs, with a minimum of 6 replicates run per treatment.

### 2.5 Coupled GC-mass spectrometry (GC-MS) analysis

Coupled GC-mass spectrometry (GC-MS) analysis was performed for the tentative identification of electrophysiologically active peaks, using the same representative samples tested in GC-EAG analysis. VOC extracts (4 µL) were analysed on an Agilent 5977B GC-MSD fitted with a non-polar HP1 column (50 m length × 0.32 mm inner diameter × 0.52 µm film thickness, J&W Scientific), using the following conditions: 30°C for 5 mins, rising 5°C min^-1^ to 150°C followed by a 10°C min^-1^ rise to 230°C for a total run time of 60 min. Ionization was by electron impact (70 eV, 220°C). Tentative identification of compounds was achieved by comparison of spectra with the NIST mass spectral library (2020, NIST, Gaithersburg, USA). Identification of compounds was confirmed by GC peak enhancement via co-injection with authentic standards,^34^ using an Agilent 6890A GC fitted with a non-polar HP1 column (50 m length × 0.32 mm inner diameter × 0.52 µm film thickness, J&W Scientific) starting at an oven temperature of 30°C for 5 mins, rising 5°C min^-1^ to 150°C, followed by a 10°C min^-1^ rise to 230°C for a total run time of 60 min. Quantification of electrophysiologically active compounds was determined from GC and GC-MS datasets using known amounts of alkane standards and dried shoot weights (Table S1).^35^

### 2.6 Chemicals

Ethylbenzene (99%), cyclohexanone (99%), heptanal (95%), benzaldehyde (99%), hexanoic acid (98%), 6-methyl-5-hepten-2-one (99%), nonanoic acid (96%), octanal (99%), nonanal (95%), undecane (97%), decanal (99%), undecanal (97%), 4-ethylbenzoic acid (99%), tetradecane (99%), hexadecane (99%) and heptadecane (99%) were obtained from Sigma-Aldrich (Gillingham, Dorset, UK). Acetoxyacetone (97%) and 3-ethylphenol (95%) were obtained from Thermo Fisher Scientific (Hemel Hempstead, Hertfordshire, UK), 3-ethylbenzaldehyde (95%) obtained from Flourochem Ltd (Hadfield, Derbyshire, UK) and pentadecane (99%) obtained from Koch-Light Laboratories Ltd (Colnbrook, Buckinghamshire, UK).

(*E*)-β-Farnesene was synthesised in-house using a modified route as previously reported. ^36^ To a solution of (*E,E*)-farnesol (1 g, 4.48 mmol) and 3,4-dihydropyran (1.88 g, 22.40 mmol) in dichloromethane (DCM; 30 ml), under nitrogen, was added *p*-toluenesulphonic acid (82 mg, 0.45 mmol) and the reaction stirred for 60 mins. The reaction mixture was diluted with DCM before being washed with water, sat NaHCO3, dried (MgSO4) and concentrated under vacuum. The crude product was purified on silica gel (4% diethyl ether in petroleum ether) to give 2-(((*2E, 6E*)-3,7,11- trimethyldodeca-2,6,10-trien-1-yl)oxy)tetrahydro-*2H*-pyran (1.00 g, 73% yield) as a colourless oil. To a solution of 2-(((*2E, 6E*)-3,7,11-trimethyldodeca-2,6,10-trien-1-yl)oxy)tetrahydro-*2H*-pyran (500 mg, 1.63 mmol) and 18-crown-6 (87 mg, 0.33 mmol) in tetrahydrofuran (THF; 25 mL), under nitrogen, was added potassium *t*-butoxide (1.83 g, 16.30 mmol) and the reaction mixture heated to 65°C for 8 h. The reaction mixture was poured into water and extracted with petroleum ether. The combined organics were dried (MgSO4) and concentrated under vacuum. The crude product was purified on silica gel (100% petroleum ether) to give (*E*)-7,11-dimethyl-3-methylenedodeca-1,6,10-triene ((*E*)-β-farnesene; 232 mg, 70% yield) as a colourless oil. All spectroscopic data was consistent with previously reported literature values.

### 2.7 Statistical analysis

Statistical analysis of olfactometry data was carried out by comparing mean time alate aphids spend between treatment and control arms (fixed effect) using a linear mixed model (LMM) in GenStat® (2016, 18th Edition, ©VSN International Ltd, Hemel Hempstead, UK), accounting for random effects caused by replicates and olfactometer area. Data were visualised using R.^37^

For the statistical comparison of the concentrations of EAG active compounds between treatments, a dataset showing the standardised amount (ng g^-1^ leaf weight hr^-1^) for the identified EAG-active compounds in each VOC extract sample across all treatments was created. Standardisation was carried out using the collected dried shoot weights and GC peak areas from an alkane C7-C22 standard (100 ng each). Using R, the dataset was log2 transformed and comparison between treatments was carried out via principle component analysis (PCA) and adonis permutational multivariate analysis test using the packages ‘vegan’, ‘ropls’ and ‘factoextra’.^38–40^

## 3 Results

### 3.1 Olfactometry assays

Alate (winged) *S. avenae* spent less time (*P*<0.05) in the arms containing the VOCs collected over a 24 h period from *S. avenae* (n = 1, 5, 10 and 25)-damaged *T. monococcum* MDR049 compared to the solvent control (Fig 1). However, alate *S. avenae* spent less time (*P*<0.05) in the arms containing the VOCs collected over a 24 h period from *S. avenae* (n = 25)-damaged *T. monococcum* MDR045 i.e., only the highest aphid density, compared to the solvent control (Fig 1). Alate *S. avenae* spent more time (*P*<0.05) in the arms containing the VOCs from *S. avenae* (n = 10, 25)-damaged *T. aestivum* var Solstice and VOCs from *S. avenae* (n = 1, 10)-damaged *T. monococcum* MDR037 compared to the solvent controls (Fig 1.)

**Figure 1.**
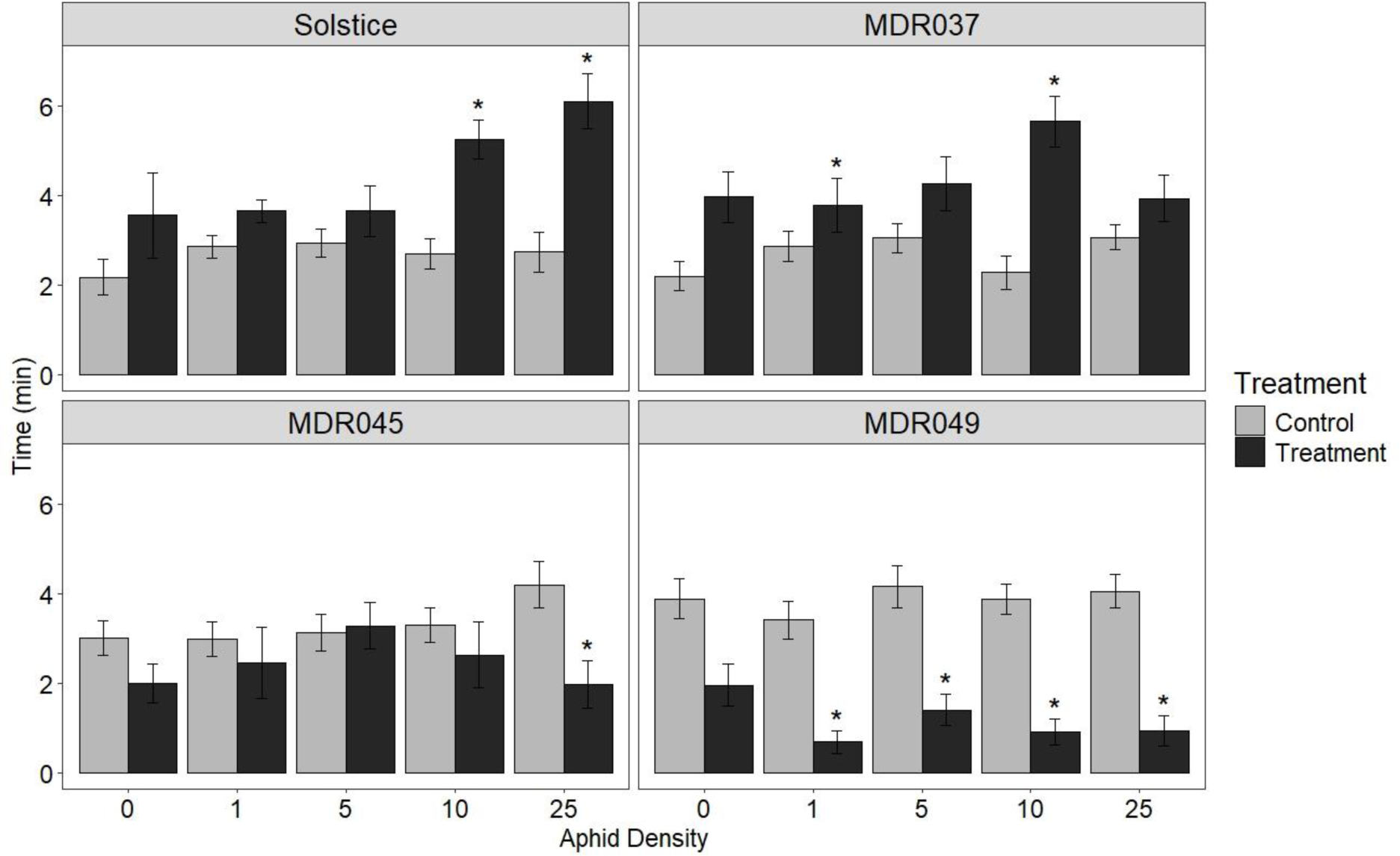
Behavioural response of alate grain aphids, *Sitobion avenae*, in a four-arm olfactometer to VOCs collected for a period of 24 h from *Triticum aestivum* var Solstice and *Triticum monococcum* MDR037, MDR045 and MDR049 plants. VOCs were collected from plants with different aphid densities (n = 0, 1, 5, 10, 25 per plant). Data are presented as the mean (min±SE) time spent in treatment and control olfactometer arms. The control in all treatments was diethyl ether. Asterisks indicate a significant difference between treatment and control (LMM test: P<0.05).

Alate *S. avenae* spent less time (mean time = 1.25 min ± 0.27, P<0.001) in the arms containing the VOCs collected over a 24 h period from *S. avenae* (n = 25)-damaged *T. monococcum* MDR049 compared to the solvent control (mean time = 5.49 min ± 0.69) in an olfactometer repellence test (Fig. S1).

### 3.2 Electrophysiology and GC-MS analysis

Coupled GC-EAG analysis using the antennae of alate *S. avenae* located 24 electrophysiologically active compounds (Table 1) from VOC extracts collected from *T. monococcum* MDR037, MDR045, MDR049 and *T. aestivum* Solstice under 0 and 25 *S. avenae* infestation densities (Fig. 2 and Fig. S2). Coupled GC-MS analysis and GC peak enhancement with authentic standards identified 21 of the 24 electrophysiologically active compounds as acetoxyacetone, ethylbenzene, cyclohexanone, heptanal, benzaldehyde, hexanoic acid, 6-methyl-5-hepten-2-one, octanal, nonanal, undecane, 3-ethylbenzaldehyde, 3-ethylphenol, decanal, nonanoic acid, undecanal, 4-ethylbenzoic acid, tetradecane, (*E*)-β-farnesene, pentadecane, hexadecane and heptadecane (Table 1). 3,4- Dimethylbenzaldehyde, 2-isopropyl-5-methyl-1-heptanol and 1-ethyldecylbenzene identity could not be confirmed as authentic standards were not commercially available at the time of experiments.

**Figure 2.**
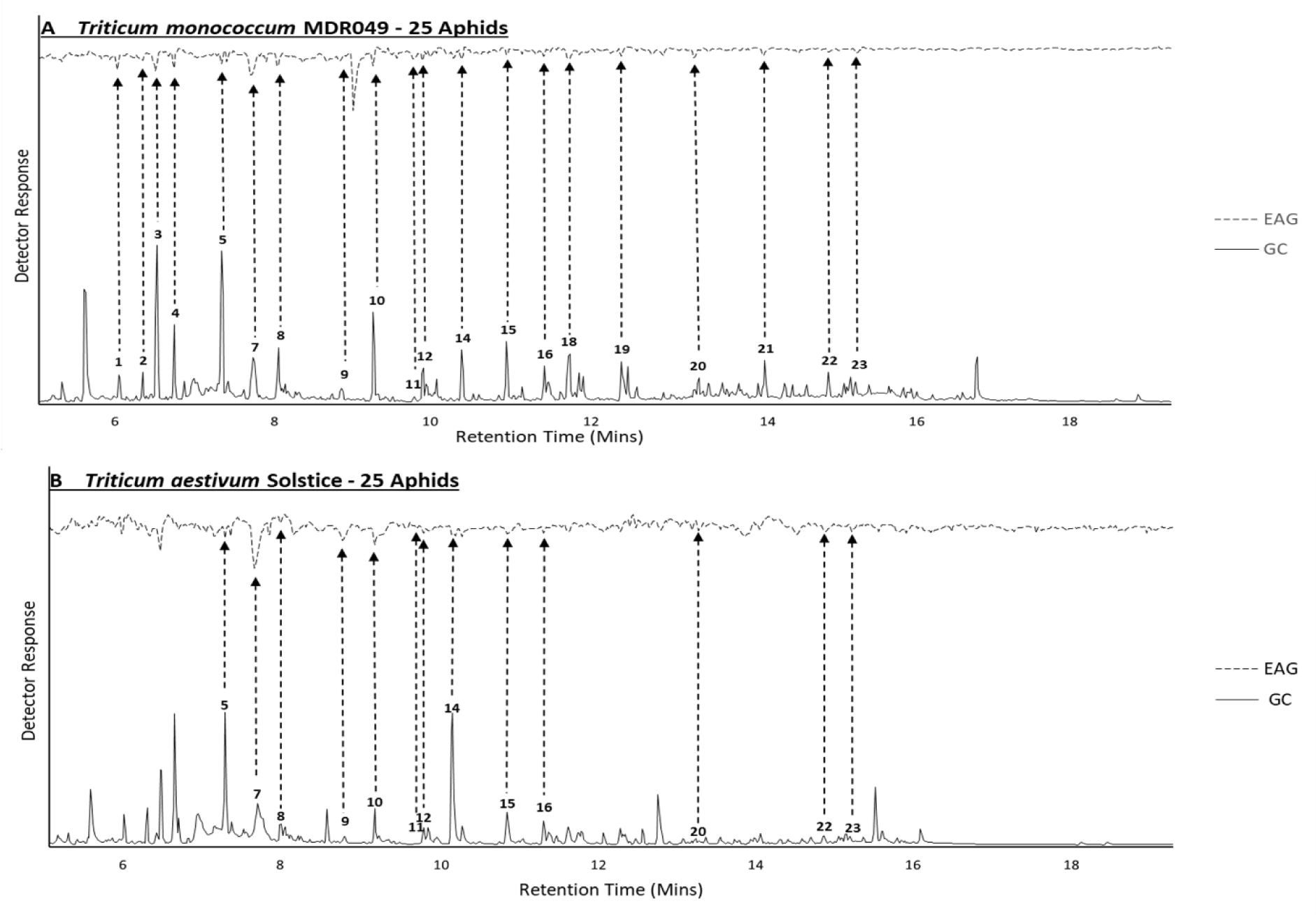
Representative coupled GC-EAG traces showing antennal responses of alate grain aphids, *Sitobion avenae* against headspace extracts collected from *S. avenae* (n=25)-infested *Triticum monococcum* MDR049 (Fig 2A) and *T. aestivum* Solstice (Fig 2B). Upper trace, response of antenna; lower trace, FID response. GC peak numbers correspond to compounds listed in Table 1 with arrows indicating their respective EAG peak. Identifications confirmed by GC peak enhancement using authentic standards.

**Table 1.**
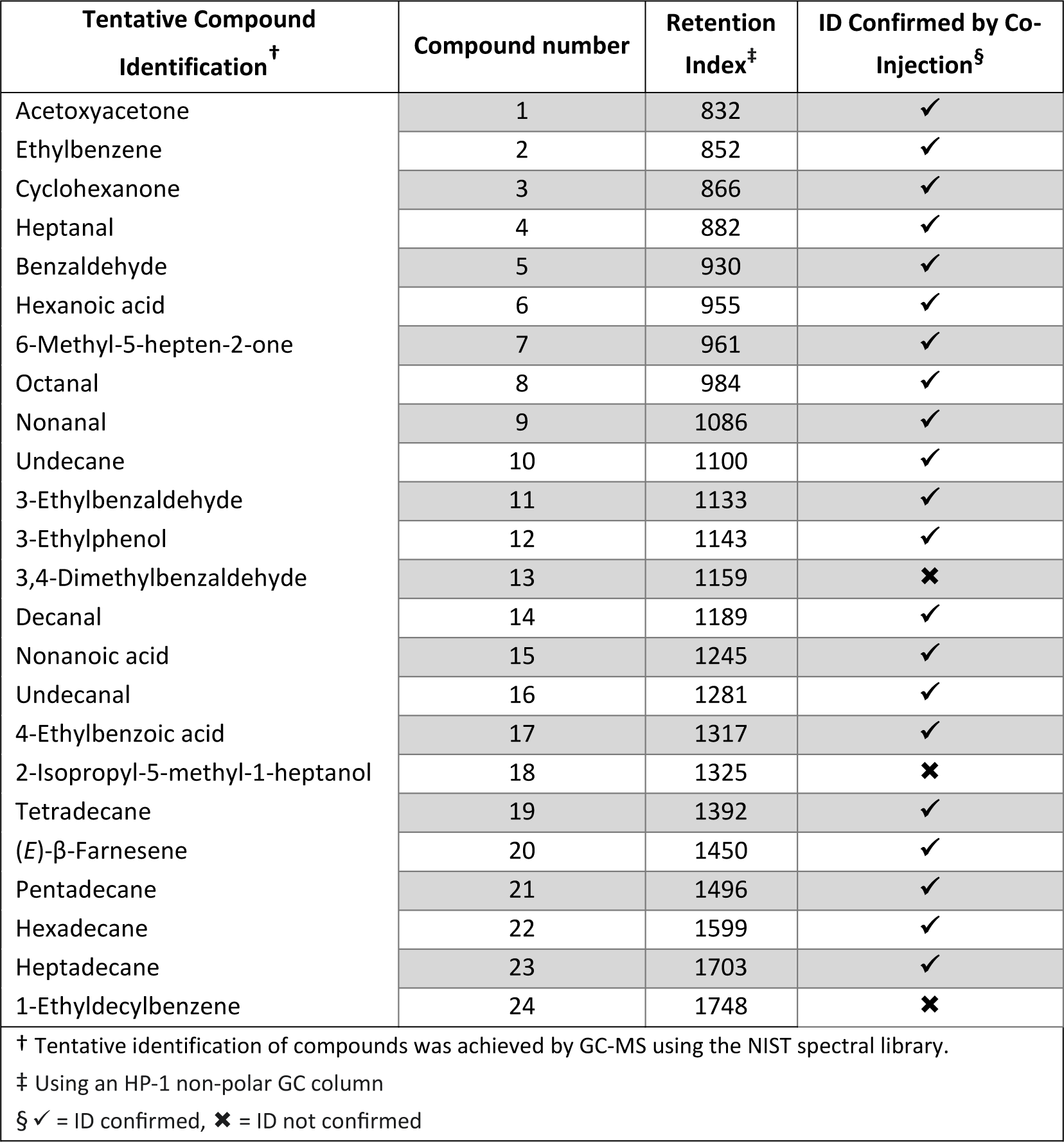
Electrophysiologically active compounds against *Sitobion avenae* alate antennae in headspace extracts of *Triticum monoccocum* MDR049, MDR045, MDR037 and *T. aestivum* Solstice collected under 0 and 25 *S. avenae* infestation densities, showing their tentative identification, assigned compound number, retention index (Kováts index) and ID confirmation by GC co-injection (✓ = ID confirmed, ® = ID not confirmed).

PCA and multivariate analysis showed no significant differences in the VOC extract composition of the 21 identified electrophysiologically active compounds between *T. monococcum* MDR037, MDR045, MDR049 and *T. aestivum* var Solstice across *S. avenae* density treatments (n = 0, 1, 5, 10, 25) (Fig. S3).

### 3.3 Behavioural activity of synthetic blends

In four-arm olfactometer bioassays, alate *S. avenae* spent less time (P<0.05) in the arms containing synthetic blends of electrophysiologically active compounds identified from the VOC extracts of MDR045 and MDR049 compared to solvent controls (Fig 3), when presented at a dose equivalent to an n = 25 and n = 5, 10, 25 aphid density for MDR045 and MDR049, respectively. Alate *S. avenae* spent more time (*P*<0.05) in the arms containing synthetic blends of electrophysiologically active compounds identified from the VOC extracts of *T. aestivum* var Solstice and MDR037 compared to solvent controls (Fig 3), when presented at a dose equivalent to an n = 10, 25 and n = 10 aphid density for Solstice and MDR037 respectively.

**Figure 3.**
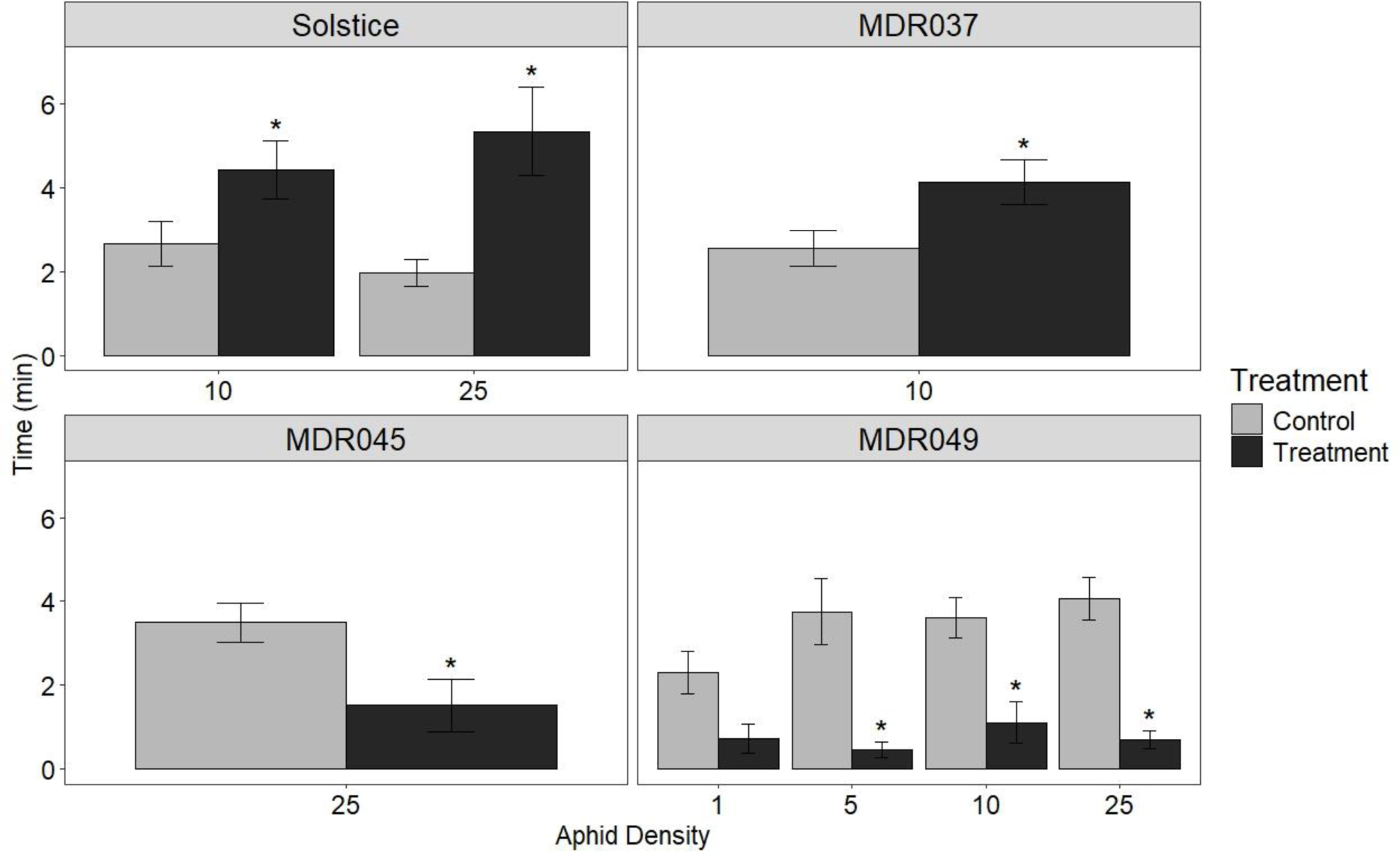
Behavioural response of alate grain aphids, *Sitobion avenae*, in a four-arm olfactometer to synthetic blends of electrophysiologically active compounds identified from *Triticum aestivum* var Solstice and *Triticum monococcum* MDR037, MDR045 and MDR049 plants which induced a significant behavioural response in Figure 1. Data are presented as the mean (min±SE) time spent in treatment and control olfactometer arms. The control in all treatments was diethyl ether. Asterisks indicate a significant difference between treatment and control (LMM test: P<0.05).

## 4 Discussion

Our study provides a mechanistic understanding for aphid antixenosis in ancestor wheat, MDR045 and MDR049, where previous field observations showed reduced *R. padi* and *S. avenae* numbers on both accessions.^18^ In olfactometry assays, VOC extracts from MDR049 induced significant reduced preference in alate *S. avenae,* compared to controls, starting from an *S. avenae* density of n=1. In MDR045, VOC extracts induced significant reduced preference in alate *S. avenae*, compared to controls, from an *S. avenae* density of n=25. This suggests that MDR049 has a threshold of one aphid required to induce VOC-related resistance mechanisms, with MDR045 having a 25-aphid threshold to induce a similar resistance mechanism. This reduced aphid preference is observed to be aphid species-specific, previous work showing a significantly reduced *R. padi* preference compared to controls in olfactometry assays by MDR049 VOC extracts collected under a 50 *R. padi* infestation density (showing similar results to those observed here against *S. avenae*).^22^ However, MDR045 VOC extracts collected under a 50 *R. padi* infestation density induced no change in preference against *R. padi* compared to controls,^22^ contrasting the reduced preference from aphid-infested MDR045 VOC extracts against *S. avenae* shown here. This species-specific response has been previously observed in wheat, where VOC extracts from primed plants induced different responses in *R. padi* and *S. avenae* in olfactometry assays,^41^ highlighting the need to test against multiple aphid species when screening for aphid resistance in crops.

Similar reduced preference of *S. avenae* has been observed in *T. monococcum* however, the cultivar tested was not provided by the authors.^42^ Contrastingly, choice assays have identified the Brazilian cultivar *T. aestivum* BRS Timbauva, bred for BYDV resistance, as being disruptive to *R. padi* behaviour.^12^ Headspace extracts from uninfested *T. aestivum* Ciko induced a preferential response in *R. padi;*^43^ however, under aphid infestation, headspace extracts became repellent.^44^ This variability is not limited to wheat but is also observed in *Zea mays* (L.).^45^ These studies highlight that VOC-mediated aphid resistance mechanisms are plant genotype-dependent and can vary highly between cultivars, explaining the variability in headspace extract bioactivity across *T. monococcum* MDR lines.

MDR049 VOC extracts induced a significant reduced preference to alate *S. avenae* at a 1 *S. avenae* density, with MDR045 VOC extracts inducing this response at a 25 *S. avenae* density, compared to controls. The results showed a lack of a gradual reduction in *S. avenae* preference with increasing aphid densities within the two accessions. MDR049 VOC extracts collected at a 0 aphid density already induced low preference towards alate *S. avenae*, compared to controls; therefore, a gradual decrease in *S. avenae* preference was unlikely since the initial preference was already low. In fact, aphid densities higher than 25 aphids, i.e. at 30, 70 and 100 aphid densities, induce distinct VOC composition changes at each density treatment in *Arabidposis thaliana* ((L.) Heynh.) under *Myzus persicae* (Sulzer) feeding.^46^ A previous study investigating the effects of varying aphid densities on plant resistance has shown that the induction of the phytohormones salicylic acid (SA) and jasmonic acid (JA), both responsible for aphid resistance induction,^47^ increases in an aphid density-dependent manner in *Medicago truncatula* (Gaertn.).^48^ In the case of *T. monococcum*, aphid density was shown to be positively correlated with presence of aphid predators on MDR045 and MDR049 in the field,^18^ which was also observed on *Rubus idae*us (L.).^49^ The current study shows that the aphid threshold required to induce resistance is genotype-dependent. Aphid density-dependent effects have also been demostrated to be species-specific, i.e. the aphid density required to induce significant yield losses in winter wheat was a minimum of 10 aphids per plant for *R. padi* and *D. noxia,* and 15 aphids per plant for *Schizaphis graminum* (Rondani).^50^

No unique EAG-active compounds were identified in either the resistant or susceptible accessions, indicating the ratio of these compounds is the key factor in inducing the different behaviroual responses observed in the olfactometry assays. Interestingly, several compounds detected as EAG-active in headspace extracts from certain accessions did not elicit a response in others, despite being present in the VOC blend. It has been shown that EAG responses are dose-dependent, so the concentration of EAG-active VOCs in headspace extracts where they did not elicit a response may have been too low to be detected by the aphid antennae.^51^ No significant differences were observed in the composition of headspace extracts between accessions when electrophysiologically active compounds were considered. This indicates that statistically non-significant but biologically important changes in VOC ratios are responsible for the contrasting behavioural activity induced by the aphid-susceptible (Solstice and MDR037) and resistant (MDR045 and MDR049) accessions. This was confirmed by testing synthetic blends of the identified electrophysiologically active VOCs at their natural doses, which induced the same behavioural response as their corresponding headspace extracts.

The importance of compound ratios in VOC blend activity against aphids has been previously demonstrated.^52^ For example, for the black-bean aphid, *Aphis fabae* (Scopoli), a blend of ten compounds from headspace extracts of its host plant were identified as being responsible for eliciting behavioural preference, whilst the individual compounds themselves induced an antixenosis response.^53–55^ Similarly, a synthetic blend of 32:1 (*E*)-2-hexenal: (*E*)-caryophyllene that simulated the natural proportions found in *Humulus lupulus* (L.) headspace attracted *Phorodon humuli* (Schrank); however, this preference was lost at a 1:1 ratio.^56^ It has been suggested that blends of active VOCs are perceived as distinct odours and therefore when in contact with individual components of the blend or with blends at different ratios, the odour becomes unrecognisable to the insect, explaining this change in behavioural response.^57^ Due to plant-plant variation, VOC blends of the same genotype may differ when grown under the same conditions,^58^ and it has been proposed that a major compound in a VOC blend can be replaced by other compounds to elicit the same response, accounting for this natural variation.^52^ It was not determined in the current study which compounds are crucial for the observed behavioural activity (antixenosis via repellent compounds and preference by attractants); thus, further work is required to test synthetic blends in olfactometry assays omitting single compounds to check if the observed responses change. Furthermore, this study assessed VOC extracts collected from 14-day old plants, but it is unknown whether antixenosis is maintained at different stages of plant growth in MDR045 and MDR049. VOC profiles have been shown to change across different plant growth stages,^59,60^ which in turn could affect VOC bioactivity. Whether or not MDR045 and MDR049 maintain antixenosis at different plant growth stages should be investigated in the future.

Of the 21 identified EAG-active compounds, 12 are green leaf volatiles (GLVs) and related compounds: heptanal, octanal, nonanal, decanal, undecanal, hexanoic acid, nonanoic acid, undecane, tetradecane, pentadecane, hexadecane and heptadecane. GLVs are common plant VOCs known to be involved in biotic resistance responses against both herbivory and pathogen infection.^61–64^ This is also the case for benzenoids,^43,53,65^ forming the second largest class of active compounds identified in *T. monococcum* headspace extracts, including ethylbenzene, benzaldehyde, 3-ethylbenzaldehyde, 3- ethylphenol, 3,4-dimethylbenzaldehyde, 4-ethylbenzoic acid and 1-ethyldecylbenzene. Many of the identified compounds have been documented to be physiologically active against aphids and involved in both attractive/preferential and repellent/antixenotic responses, depending on their ratios in blends. Heptanal, nonanal, decanal, benzaldehyde, tetradecane, hexadecane, heptadecane and 4- ethylbenzaldehyde were identified as active compounds, using olfactometry assays, in headspace extracts from wheat and oat, inducing preference in *R. padi* alate.^43^ When tested individually, only heptanal, nonanal, decanal and benzaldehyde induced preference. Similarly, octanal, decanal, undecanal, (*E*)-β-farnesene, benzaldehyde and 6-methyl-5-hepten-2-one were identified as physiologically active compounds from field bean *Vicia faba* L., headspace extracts that induced preference in *A. fabae.*^53^ When tested individually, octanal, decanal, undecanal and (*E*)-β-farnesene induced an antixenotic response, whilst benzaldehyde, undecanal and 6-methyl-5-hepten-2-one were inactive; however, preference could be induced when they were combined in their naturally occurring ratios.^54^

The VOCs identified in this study also play a role in tri-trophic cereal-aphid-virus interactions. Nonanal, decanal and undecane are present in headspace extracts from BYDV viruliferous and non-viruliferous *T. aestivum* Lambert, with the individual compounds and BYDV-infected headspace extracts eliciting a higher preference in *R. padi* in immigration bioassays, compared to the solvent control.^66^ Decanal concentrations in VOC blends of BYDV viruliferous and non-viruliferous *T. aestivum* Aikang-58 were shown to be important for the preferential response of BYDV viruliferous and non-viruliferous *S. graminum*, with non-viruliferous aphids preferring viruliferous plants with higher decanal concentrations, whilst viruliferous aphids preferring non-viruliferous plants with lower decanal concentrations.^67^ Nonanoic acid, nonanal, (*E*)-β-farnesene, ethylbenzene, cyclohexanone and benzaldehyde have all previously been shown to be involved in aphid predator attraction.^68–70^ Considering the field observations of higher aphid predator presence in MDR049,^18^ the presence of these compounds in headspace extracts of MDR045 and MDR049 may indicate that part of the aphid resistance mechanisms of these accessions involves aphid predator attraction; however, this will need further laboratory testing for confirmation.

## 5 Conclusion

In conclusion, the results in this study suggest a role for VOCs in aphid antixenosis displayed by *T. monococcum* MDR045 and MDR049. The induced VOCs appear to be present across the aphid-resistant and susceptible accessions, implying that VOC blend composition is crucial for conveying the antixenosis effect. Furthermore, these results provide a platform for the incorporation of VOC-based aphid resistance into modern elite wheat varieties, which can then be deployed in IPM strategies to reduce the direct and indirect negative impacts of aphid infestations, thereby reducing dependence on insecticides. Further work is required to elucidate the functional genes involved in the production of VOCs in MDR045 and MDR049 and the mechanisms involved in their regulation.

## Acknowledgements

We would like to thank Suzanne Clark for statistical and experimental design advice. We also thank the Wheat Genetic Improvement Network (http://www.wgin.org.uk/) for providing *T. monococcum* seeds and which is supported by a grant from the UK Government Department for Environment, Food and Rural Affairs (Defra, AR0709). ANB was supported by a Biotechnology and Biological Sciences Research Council (BBSRC) studentship (2019-2023) awarded to Rothamsted Research and University of Nottingham. Rothamsted Research receives strategic funding from BBSRC. This work formed part of the Rothamsted Smart Crop Protection (SCP) strategic programme (BBS/OS/CP/000001) funded through BBSRC’s Industrial Strategy Challenge Fund. We acknowledge support from the Growing Health Institute Strategic Programme [BB/X010953/1; BBS/E/RH/230003A].

## References

1. 1. FAO. FAOSTAT. https://www.fao.org/faostat/en/#home (2023).

2. Gildow, F. E. & Rochow, W. F. Barley Yellow Dwarf in California: Vector Competence and Luteovirus Identification. Plant Dis. 67, 140–143 (1982).

3. Rabbinge, R., Drees, E. M., van der Graaf, M., Verberne, F. C. M. & Wesselo, A. Damage effects of cereal aphids in wheat. *Netherlands J*. Plant Pathol. 87, 217–232 (1981).

4. Aradottir, G. I. & Crespo-Herrera, L. Host plant resistance in wheat to barley yellow dwarf viruses and their aphid vectors: a review. Curr. Opin. Insect Sci. 45, 59–68 (2021).

5. Tanguy, S. & Dedryver, C. A. Reduced BYDV-PAV transmission by the grain aphid in a Triticum monococcum line. Eur. J. Plant Pathol. 123, 281–289 (2009).

6. Foster, S. P. et al. A mutation (L1014F) in the voltage-gated sodium channel of the grain aphid, Sitobion avenae, is associated with resistance to pyrethroid insecticides. Pest Manag. Sci. 70, 1249–1253 (2014).

7. Pickett, J. A. Food security: Intensification of agriculture is essential, for which current tools must be defended and new sustainable technologies invented. Food Energy Secur. 2, 167–173 (2013).

8. Sparks, T. C. Insecticide discovery : An evaluation and analysis. Pestic. Biochem. Physiol. 107, 8–17 (2013).

9. Tolmay, V. L., Sydenham, S. L., Sikhakhane, T. N., Nhlapho, B. N. & Tsilo, T. J. Elusive diagnostic markers for Russian wheat aphid resistance in bread wheat: Deliberating and reviewing the status quo. Int. J. Mol. Sci. 21, 1–22 (2020).

10. Walls, J., Rajotte, E. & Rosa, C. The past, present, and future of barley yellow dwarf management. Agric. 9, 1–16 (2019).

11. Zhang, Y., Fan, J., Francis, F. & Chen, J. Watery Saliva Secreted by the Grain Aphid Sitobion avenae Stimulates Aphid Resistance in Wheat. J. Agric. Food Chem. 65, 8798–8805 (2017).

12. Correa, L. D. J. et al. A Comprehensive Analysis of Wheat Resistance to Rhopalosiphum padi (Hemiptera: Aphididae) in Brazilian Wheat Cultivars. J. Econ. Entomol. 113, 1493–1503 (2020).

13. Mansouri, S. M., Norouzi, R. & Mehrparvar, M. Resistance of wild wheat, Triticum boeoticum Boiss. to Schizaphis graminum (Rondani) (Hem.: Aphididae). North. West. J. Zool. 15, 1–6 (2019).

14. Radchenko, E. E. Resistance of Triticum species to cereal aphids. Czech J. Genet. Plant Breed. 47, 2009–2012 (2011).

15. Migui, S. M. & Lamb, R. J. Patterns of resistance to three cereal aphids among wheats in the genus Triticum (Poaceae) . Bull. Entomol. Res. 93, 323–333 (2003).

16. Elek, H. et al. Aphid resistance in wheat varieties. Commun. Agric. Appl. Biol. Sci. 74, 233–241 (2009).

17. Singh, B. et al. Characterisation of bird cherry-oat aphid (Rhopalosiphum padi L.) behaviour and aphid host preference in relation to partially resistant and susceptible wheat landraces. Ann. Appl. Biol. 177, 184–194 (2020).

18. Simon, A. L., Caulfield, J. C., Hammond-Kosack, K. E., Field, L. M. & Aradottir, G. I. Identifying aphid resistance in the ancestral wheat Triticum monococcum under field conditions. Sci. Rep. 11, 1–12 (2021).

19. Simon, A. L., Wellham, P. A. D., Aradottir, G. I. & Gange, A. C. Unravelling mycorrhiza-induced wheat susceptibility to the English grain aphid Sitobion avenae. Sci. Rep. 7, 1–11 (2017).

20. Greenslade, A. F. C. et al. Triticum monococcum lines with distinct metabolic phenotypes and phloem-based partial resistance to the bird cherry-oat aphid Rhopalosiphum padi. Ann. Appl. Biol. 168, 435–449 (2016).

21. Haas, M., Schreiber, M. & Mascher, M. Domestication and crop evolution of wheat and barley: Genes, genomics, and future directions. J. Integr. Plant Biol. 61, 204–225 (2019).

22. Simon, A. L. Unravelling cereal aphid interactions with ancestral wheat Triticum monococcum. (University of Nottingham, 2020).

23. Turlings, T. C. J. & Erb, M. Tritrophic Interactions Mediated by Herbivore-Induced Plant Volatiles: Mechanisms, Ecological Relevance, and Application Potential. Annu. Rev. Entomol. 63, 433–452 (2018).

24. Moraes, M. C. B. et al. Attraction of the stink bug egg parasitoid Telenomus podisi to defence signals from soybean activated by treatment with cis-jasmone. Entomol. Exp. Appl. 131, 178– 188 (2009).

25. Zhou, S. & Jander, G. Molecular ecology of plant volatiles in interactions with insect herbivores. J. Exp. Bot. 73, 449–462 (2022).

26. Pickett, J. A. & Khan, Z. R. Plant volatile-mediated signalling and its application in agriculture: successes and challenges. New Phytol. 212, 856–870 (2016).

27. Chen, K., Wang, Y., Zhang, R., Zhang, H. & Gao, C. CRISPR/Cas Genome Editing and Precision Plant Breeding in Agriculture. Annu. Rev. Plant Biol. 70, 667–697 (2019).

28. Bruce, T. J. A. et al. The first crop plant genetically engineered to release an insect pheromone for defence. Sci. Rep. 5, 1–9 (2015).

29. Li, F. et al. Expression of lima bean terpene synthases in rice enhances recruitment of a beneficial enemy of a major rice pest. Plant Cell Environ. 41, 111–120 (2018).

30. Zhang, L. et al. Overexpression of the caryophyllene synthase gene GhTPS1 in cotton negatively affects multiple pests while attracting parasitoids. Pest Manag. Sci. 76, 1722–1730 (2020).

31. MacGillivray, M. E. & Anderson, G. B. Three Useful Insect Cages. Can. Entomol. 89, 43–46 (1957).

32. Pettersson, J. An Aphid Sex Attractant. Ent. scand. 63–73 (1970).

33. Wadhams, L. J. The Use of Coupled Gas Chromatography: Electrophysiological Techniques in the Identification of Insect Pheromones. Chromatogr. Isol. Insect Horm. Pheromones (1990).

34. Pickett, J. A. GC-MS in insect pheromone identification: three extreme case histories. Chromatogr. Isol. Insect Horm. Pheromones 299–309 (1990).

35. Kost, C. & Heil, M. Herbivore-induced plant volatiles induce an indirect defence in neighbouring plants. J. Ecol. 94, 619–628 (2006).

36. Arkoudis, E. & Stratakis, M. Synthesis of Cordiaquinones B, C, J, and K on the Basis of a Bioinspired Approach and the Revision of the Relative Stereochemistry of Cordiaquinone C. J. Org. Chem. 73, 4484–4490 (2008).

37. 37. Team, R. C. R: A language and environment for statistical computing. at https://www.r-project.org/ (2023).

38. Oksansen, J., Simspon, G., Blanchet, G., Kindt, R. & Legendre, P. vegan: Community Ecology Package. R package version 2.6–4. at https://cran.r-project.org/package=vegan (2022).

39. 39. Kassambara, A. & Mundt, F. factoextra: Extract and Visualize the Results of Multivariate Data Analyses. R package version 1.0.7. at https://cran.r-project.org/package=factoextra (2020).

40. Thévenot, E. A., Roux, A., Xu, Y., Ezan, E. & Junot, C. Analysis of the Human Adult Urinary Metabolome Variations with Age, Body Mass Index, and Gender by Implementing a Comprehensive Workflow for Univariate and OPLS Statistical Analyses. J. Proteome Res. 14, 3322–3335 (2015).

41. Pettersson, J., Quiroz, A. & Fahad, A. E. Aphid antixenosis mediated by volatiles in cereals. Acta Agric. Scand. Sect. B Soil Plant Sci. 46, 135–140 (1996).

42. Hu, X. S. et al. Resistance of wheat accessions to the english grain aphid Sitobion avenae. PLoS One 11, 1–17 (2016).

43. Quiroz, A. & Niemeyer, H. M. Olfactometer-assessed responses of aphid Rhopalosiphum padi to wheat and oat volatiles. J. Chem. Ecol. 24, 113–124 (1998).

44. Quiroz, A., Pettersson, J., Pickett, J. A., Wadhams, L. J. & Niemeyer, H. M. Semiochemicals mediating spacing behavior of bird cherry-oat aphid. J. Chem. Ecol. 23, 2599–2607 (1997).

45. Schröder, M. L., Glinwood, R., Webster, B., Ignell, R. & Krüger, K. Olfactory responses of Rhopalosiphum padi to three maize, potato, and wheat cultivars and the selection of prospective crop border plants. Entomol. Exp. Appl. 157, 241–253 (2015).

46. Truong, D. H. et al. Temperature regimes and aphid density interactions differentially influence VOC emissions in Arabidopsis. Arthropod. Plant. Interact. 8, 317–327 (2014).

47. Morkunas, I., Mai, V. C. & Gabryś, B. Phytohormonal signaling in plant responses to aphid feeding. Acta Physiol. Plant. 33, 2057–2073 (2011).

48. Stewart, S. A., Hodge, S., Bennett, M., Mansfield, J. W. & Powell, G. Aphid induction of phytohormones in Medicago truncatula is dependent upon time post-infestation, aphid density and the genotypes of both plant and insect. Arthropod. Plant. Interact. 10, 41–53 (2016).

49. Mitchell, C., Johnson, S. N., Gordon, S. C., Birch, A. N. E. & Hubbard, S. F. Combining plant resistance and a natural enemy to control Amphorophora idaei. BioControl 55, 321–327 (2010).

50. Kieckhefer, R. W. & Gellner, J. L. Yield Losses in Winter Wheat Caused by Low-Density Cereal Aphid Populations. Agron. J. 84, 180–183 (1992).

51. Birkett, M. A. et al. Electrophysiological responses of the lepidopterous stemborers Chilo partellus and Busseola fusca to volatiles from wild and cultivated host plants. J. Chem. Ecol. 32, 2475–2487 (2006).

52. Bruce, T. J. A. & Pickett, J. A. Perception of plant volatile blends by herbivorous insects - Finding the right mix. Phytochemistry 72, 1605–1611 (2011).

53. Webster, B. et al. Identification of volatile compounds used in host location by the black bean aphid, Aphis fabae. J. Chem. Ecol. 34, 1153–1161 (2008).

54. Webster, B., Bruce, T., Pickett, J. & Hardie, J. Volatiles functioning as host cues in a blend become nonhost cues when presented alone to the black bean aphid. Anim. Behav. 79, 451– 457 (2010).

55. Webster, B., Bruce, T., Pickett, J. & Hardie, J. Olfactory recognition of host plants in the absence of host-specific volatile compounds. Commun. Integr. Biol. 1, 167–169 (2008).

56. Campbell, C. A. M., Pettersson, J., Pickett, J. A., Wadhams, L. J. & Woodcock, C. M. Spring migration of damson-hop aphid, Phorodon humuli (Homoptera, Aphididae), and summer host plant-derived semiochemicals released on feeding. J. Chem. Ecol. 19, 1569–1576 (1993).

57. van Wijk, M., de Bruijn, P. J. A. & Sabelis, M. W. Complex odor from plants under attack: Herbivore’s enemies react to the whole, not its parts. PLoS One 6, (2011).

58. Webster, B., Gezan, S., Bruce, T., Hardie, J. & Pickett, J. Between plant and diurnal variation in quantities and ratios of volatile compounds emitted by Vicia faba plants. Phytochemistry 71, 81–89 (2010).

59. Kong, M., An, J., Jung, D. & Hong, T. Characterization of the aroma in barley leaves from different cultivars and tillering stages by HS-SPME/GC-MS, GC-O and E-nose. Food Biosci. (2024) doi:10.1016/j.fbio.2024.104117.

60. Azam, M., Jiang, Q., Zhang, B., Xu, C. & Chen, K. Citrus leaf volatiles as affected by developmental stage and genetic type. Int. J. Mol. Sci. 14, 17744–17766 (2013).

61. Allmann, S. et al. Feeding-induced rearrangement of green leaf volatiles reduces moth oviposition. Elife 2013, 1–23 (2013).

62. Schuman, M. C., Barthel, K. & Baldwin, I. T. Herbivory-induced volatiles function as defenses increasing fitness of the native plant Nicotiana attenuata in nature. Elife 2012, 1–29 (2012).

63. De Moraes, C. M., Lewis, W. J., Pare, P. W., Alborn, H. T. & Tumiinson, J. H. Herbivore-infested plants selectively attract parasitoids. Nature 393, 570–573 (1998).

64. Yi, H. S., Heil, M., Adame-Álvarez, R. M., Ballhorn, D. J. & Ryu, C. M. Airborne induction and priming of plant defenses against a bacterial pathogen. Plant Physiol. 151, 2152–2161 (2009).

65. Zhu, X. et al. Chemical Composition and Attractant Activity of Volatiles from Rhus potaninii to The Spring Aphid. Molecules 25, (2020).

66. Medina-Ortega, K. J., Bosque-Pérez, N. A., Ngumbi, E., Jiménez-Martínez, E. S. & Eigenbrode, S. D. Rhopalosiphum padi (Hemiptera: Aphididae) Responses to Volatile Cues From Barley Yellow Dwarf Virus–Infected Wheat . Environ. Entomol. 38, 836–845 (2009).

67. Hu, Z. et al. Barley yellow dwarf virus-infected wheat plant modulated selection behavior of vector aphids. J. Pest Sci. (2004). 95, 1273–1285 (2022).

68. Leroy, P. D., Schillings, T., Farmakidis, J. & Lognay, G. Testing semiochemicals from aphid , plant and conspecific : attraction of Harmonia axyridis. Insect Sci. 372–382 (2012) doi:10.1111/j.1744-7917.2011.01449.x.

69. Sasso, R., et al. Host-locating response by the aphid parasitoid Aphidius ervi to tomato plant volatiles. J. Plant Interact. 9145, (2008).

70. Goelen, T. et al. Identification and application of bacterial volatiles to attract a generalist aphid parasitoid : from laboratory to greenhouse assays. (2020) doi:10.1002/ps.6102.

