## Supplementary material for "Characterisation of aphid antixenosis in aphid-resistant ancestor wheat, *Triticum monococcum*"

**Table S1**. Mean normalised amount (ng $g^{-1}$ leaf weight $hr^{-1}$) of EAG-active compounds emitted by *Triticum aestivum* Solstice and *T. monococcum* MDR037, MDR045 and MDR049 plants subjected to alate *Sitobion avenae* densities (n = 0, 1, 5, 10, 25).

| *Accession* | *Aphid Density* | *Acetoxy acetone* | *Ethylbenzene* | *Cyclohexanone* | *Heptanal* | *Benzaldehyde* | *Hexanoic Acid* | *6-Methyl-5-hepten-2-one* | *Octanal* | *Nonanal* | *Undecane* | *3-EthylBenzaldehyde* | *3-EthylPhenol* | *Decanal* | *Nonanoic Acid* | *4-EthylBenzoic Acid* | *Tetradecane* | *(E)-β-Farnesene* | *Pentadecane* | *Hexadecane* | *Heptadecane* |
|---|---|---|---|---|---|---|---|---|---|---|---|---|---|---|---|---|---|---|---|---|---|
| *MDR049* | 0 | 3.2 | 9.4 | 35.3 | 28.8 | 7.3 | 2.4 | 0.7 | 6.7 | 9.2 | 0.4 | 2.2 | 0.3 | 5.7 | 0.2 | 0.4 | 3.9 | 0.4 | 0.5 | 1.8 | 1.4 |
| *MDR049* | 1 | 5.9 | 14.7 | 44.0 | 39.3 | 11.7 | 3.7 | 0.9 | 6.6 | 12.6 | 1.6 | 1.2 | 0.0 | 7.8 | 2.6 | 0.2 | 4.1 | 0.2 | 0.5 | 1.5 | 0.6 |
| *MDR049* | 5 | 5.3 | 14.5 | 31.5 | 34.6 | 8.4 | 2.9 | 0.3 | 7.9 | 13.1 | 0.4 | 0.9 | 0.2 | 7.5 | 0.6 | 0.4 | 3.8 | 0.2 | 1.3 | 1.5 | 0.9 |
| *MDR049* | 10 | 2.2 | 4.3 | 14.5 | 14.6 | 2.7 | 0.6 | 0.2 | 2.6 | 3.9 | 0.1 | 0.4 | 0.0 | 1.6 | 0.1 | 0.0 | 1.2 | 0.2 | 0.1 | 0.8 | 0.6 |
| *MDR049* | 25 | 3.8 | 9.8 | 29.7 | 23.2 | 7.4 | 1.8 | 0.0 | 6.4 | 8.8 | 0.4 | 2.8 | 0.0 | 4.0 | 0.8 | 0.1 | 2.9 | 0.6 | 0.2 | 2.0 | 1.5 |
| *MDR045* | 0 | 5.2 | 31.9 | 26.1 | 51.8 | 18.3 | 7.5 | 1.6 | 17.3 | 21.8 | 1.1 | 1.4 | 1.6 | 13.5 | 0.6 | 0.9 | 7.8 | 0.1 | 0.1 | 2.8 | 1.6 |
| *MDR045* | 1 | 3.2 | 10.5 | 21.6 | 24.3 | 6.1 | 1.8 | 0.3 | 5.4 | 9.9 | 0.4 | 1.4 | 0.3 | 5.7 | 0.4 | 0.4 | 4.0 | 0.6 | 1.7 | 2.2 | 1.6 |
| *MDR045* | 5 | 2.2 | 6.1 | 16.5 | 14.8 | 3.2 | 0.7 | 0.2 | 2.4 | 4.6 | 0.4 | 1.3 | 0.0 | 1.7 | 0.7 | 0.0 | 1.4 | 0.2 | 0.1 | 1.2 | 0.6 |
| *MDR045* | 10 | 5.7 | 14.7 | 30.6 | 37.9 | 8.9 | 2.4 | 1.1 | 8.4 | 11.3 | 0.5 | 0.8 | 0.3 | 6.2 | 0.9 | 0.3 | 2.9 | 0.1 | 0.5 | 2.2 | 0.7 |
| *MDR045* | 25 | 2.8 | 7.5 | 30.1 | 22.2 | 6.1 | 1.0 | 0.3 | 5.3 | 9.5 | 0.3 | 1.1 | 0.0 | 4.2 | 0.0 | 0.3 | 3.2 | 1.2 | 0.4 | 1.2 | 0.9 |
| *MDR037* | 0 | 3.3 | 21.1 | 51.7 | 45.9 | 13.0 | 4.5 | 0.1 | 11.8 | 16.2 | 0.7 | 1.4 | 0.2 | 10.2 | 0.7 | 0.3 | 5.1 | 0.4 | 0.3 | 3.4 | 1.7 |
| *MDR037* | 1 | 3.0 | 17.5 | 19.9 | 34.4 | 12.1 | 5.6 | 2.5 | 13.9 | 18.3 | 1.0 | 1.6 | 0.7 | 10.4 | 0.8 | 0.3 | 5.8 | 0.2 | 0.0 | 1.6 | 0.5 |
| *MDR037* | 5 | 4.2 | 19.4 | 21.9 | 41.3 | 13.8 | 5.0 | 0.4 | 12.7 | 16.5 | 0.8 | 1.7 | 0.9 | 9.3 | 0.8 | 0.4 | 5.5 | 0.5 | 0.0 | 2.8 | 1.7 |
| *MDR037* | 10 | 6.3 | 15.2 | 33.5 | 45.1 | 9.0 | 2.0 | 1.8 | 8.9 | 13.0 | 0.6 | 1.0 | 0.2 | 7.6 | 0.6 | 0.7 | 4.8 | 0.4 | 0.7 | 1.9 | 1.2 |
| *MDR037* | 25 | 3.5 | 7.9 | 18.6 | 23.4 | 5.6 | 1.6 | 0.0 | 5.7 | 8.4 | 0.2 | 0.6 | 0.1 | 3.9 | 0.0 | 0.2 | 1.1 | 1.7 | 0.1 | 0.9 | 1.0 |
| *Solstice* | 0 | 1.3 | 3.1 | 8.9 | 9.0 | 2.5 | 0.4 | 0.1 | 2.2 | 2.7 | 0.1 | 0.5 | 0.0 | 1.2 | 0.1 | 0.1 | 0.2 | 0.0 | 0.1 | 0.5 | 0.2 |
| *Solstice* | 1 | 4.4 | 30.8 | 9.5 | 56.7 | 12.8 | 6.7 | 4.8 | 14.1 | 19.9 | 1.6 | 1.7 | 0.0 | 13.2 | 0.9 | 1.1 | 10.0 | 0.4 | 0.0 | 4.9 | 3.5 |
| *Solstice* | 5 | 3.7 | 10.0 | 8.5 | 30.2 | 5.6 | 1.2 | 1.3 | 5.1 | 6.8 | 0.4 | 0.8 | 0.0 | 3.9 | 0.2 | 0.3 | 2.6 | 0.3 | 0.4 | 1.5 | 0.9 |
| *Solstice* | 10 | 2.1 | 4.9 | 11.2 | 15.5 | 6.2 | 2.0 | 0.1 | 6.0 | 10.0 | 0.4 | 1.8 | 0.5 | 6.0 | 0.7 | 3.0 | 3.8 | 0.2 | 0.3 | 1.7 | 0.7 |
| *Solstice* | 25 | 5.4 | 11.3 | 24.8 | 47.0 | 9.1 | 2.0 | 0.1 | 8.2 | 10.8 | 0.5 | 1.2 | 0.1 | 6.6 | 0.3 | 0.3 | 4.1 | 2.1 | 0.2 | 1.3 | 0.8 |

**Table S2.** Volume (µl) of authentic standard solutions (1mg/ml in diethyl ether) required to create a 10x concentration synthetic blend in diethyl ether (1 mL) for the EAG-active compounds from *Triticum aestivum* Solstice and *T. monococcum* MDR037, MDR045 and MDR049 headspace extract. Synthetic blends were subsequently diluted for use in olfactometry assays.

| *Accession* | *MDR049* | *MDR049* | *MDR049* | *MDR049* | *MDR045* | *MDR037* | *Solstice* | *Solstice* |
|---|---|---|---|---|---|---|---|---|
| *Aphid Density* | *1* | *5* | *10* | *25* | *25* | *10* | *10* | *25* |
| *Acetoxyacetone* | 4.1 | 3.4 | 1.8 | 1.8 | 1.8 | 4.8 | 1.4 | 4.1 |
| *Ethylbenzene* | 10.1 | 9.1 | 3.7 | 4.4 | 3.7 | 11.5 | 2.8 | 8.3 |
| *Cyclohexanone* | 30 | 21.1 | 12.5 | 13.6 | 17.9 | 26.1 | 8.3 | 17.1 |
| *Heptanal* | 7.8 | 5.2 | 2.6 | 3.5 | 3.6 | 6.8 | 3.9 | 7.2 |
| *Benzaldehyde* | 28 | 23.3 | 12.3 | 11.4 | 13.7 | 35.6 | 11.5 | 35.1 |
| *Hexanoic Acid* | 2.7 | 1.8 | 0.6 | 0.9 | 0.7 | 1.4 | 1.4 | 1.7 |
| *6-Methyl-5-hepten-2-one* | 0.5 | 0.2 | 0.2 | 0 | 0.2 | 1.3 | 0.1 | 0.2 |
| *Octanal* | 4.8 | 5.1 | 2.5 | 3.1 | 3.3 | 7 | 4.1 | 7 |
| *Nonanal* | 10.4 | 10 | 4.2 | 4.9 | 6.8 | 12.1 | 8.6 | 10.9 |
| *Undecane* | 1.2 | 0.3 | 0.2 | 0.2 | 0.2 | 0.5 | 0.3 | 0.5 |
| *3-Ethylbenzaldehyde* | 0.9 | 0.7 | 0.6 | 1.5 | 0.9 | 1 | 1.6 | 1.2 |
| *3-Ethylphenol* | 0 | 0.2 | 0 | 0 | 0 | 0.2 | 0.4 | 0 |
| *Decanal* | 6.4 | 5.5 | 1.8 | 2.2 | 3 | 6.6 | 4.9 | 6.3 |
| *Nonanoic Acid* | 2 | 0.5 | 0.1 | 0.4 | 0 | 0.5 | 0.8 | 0.3 |
| *4-Ethylbenzoic Acid* | 0.2 | 0.3 | 0 | 0.1 | 0.2 | 0.5 | 4.5 | 0.2 |
| *Tetradecane* | 2.9 | 2.4 | 1.1 | 1.4 | 1.9 | 3.9 | 2.4 | 3.2 |
| *(E)-β-Farnesene* | 0.1 | 0.1 | 0.2 | 0.3 | 0.8 | 0.3 | 0.1 | 1.7 |
| *Pentadecane* | 0.4 | 0.8 | 0.1 | 0.1 | 0.3 | 0.6 | 0.1 | 0.2 |
| *Hexadecane* | 1 | 1.1 | 0.7 | 1 | 0.8 | 1.6 | 1.5 | 1.1 |
| *Heptadecane* | 0.4 | 0.6 | 0.5 | 0.7 | 0.6 | 1 | 0.5 | 0.7 |

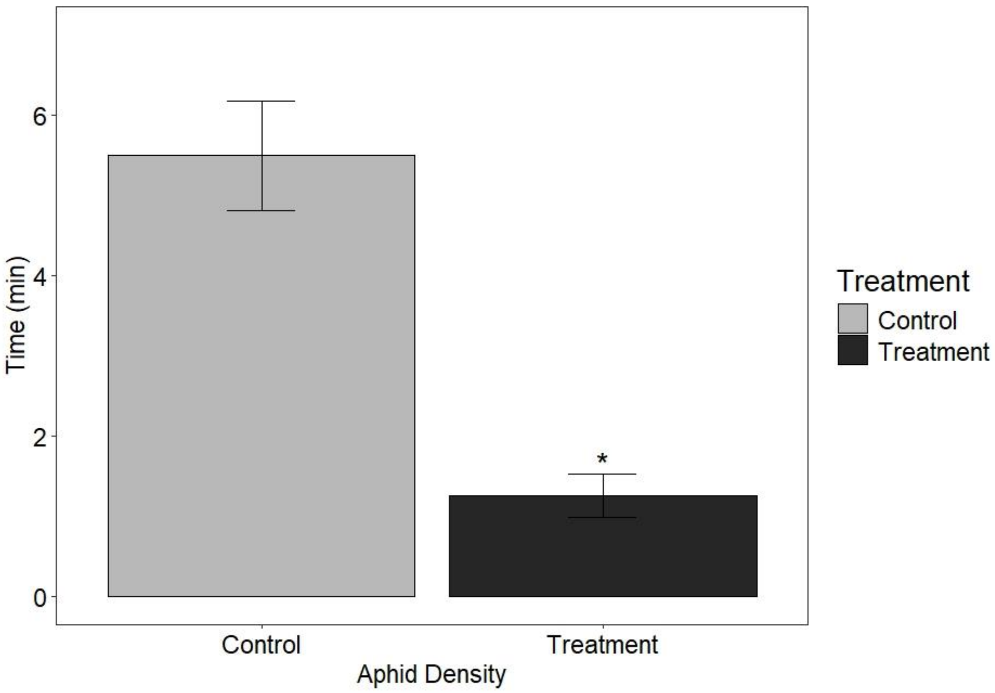


**Figure S1** Behavioural response of alate grain aphids, *Sitobion avenae*, in a four-arm olfactometer to VOCs collected for a period of 24h from *S. avenae* (n=25)-infested *Triticum monococcum* MDR049 plants. Data are presented as the mean (min ±SE) time spent in treatment and control olfactometer arms. The experiment included three treatment arms and one control arm. The control was diethyl ether. Asterisks indicate a significant difference between treatment and control (ANOVA: $P<0.05$).

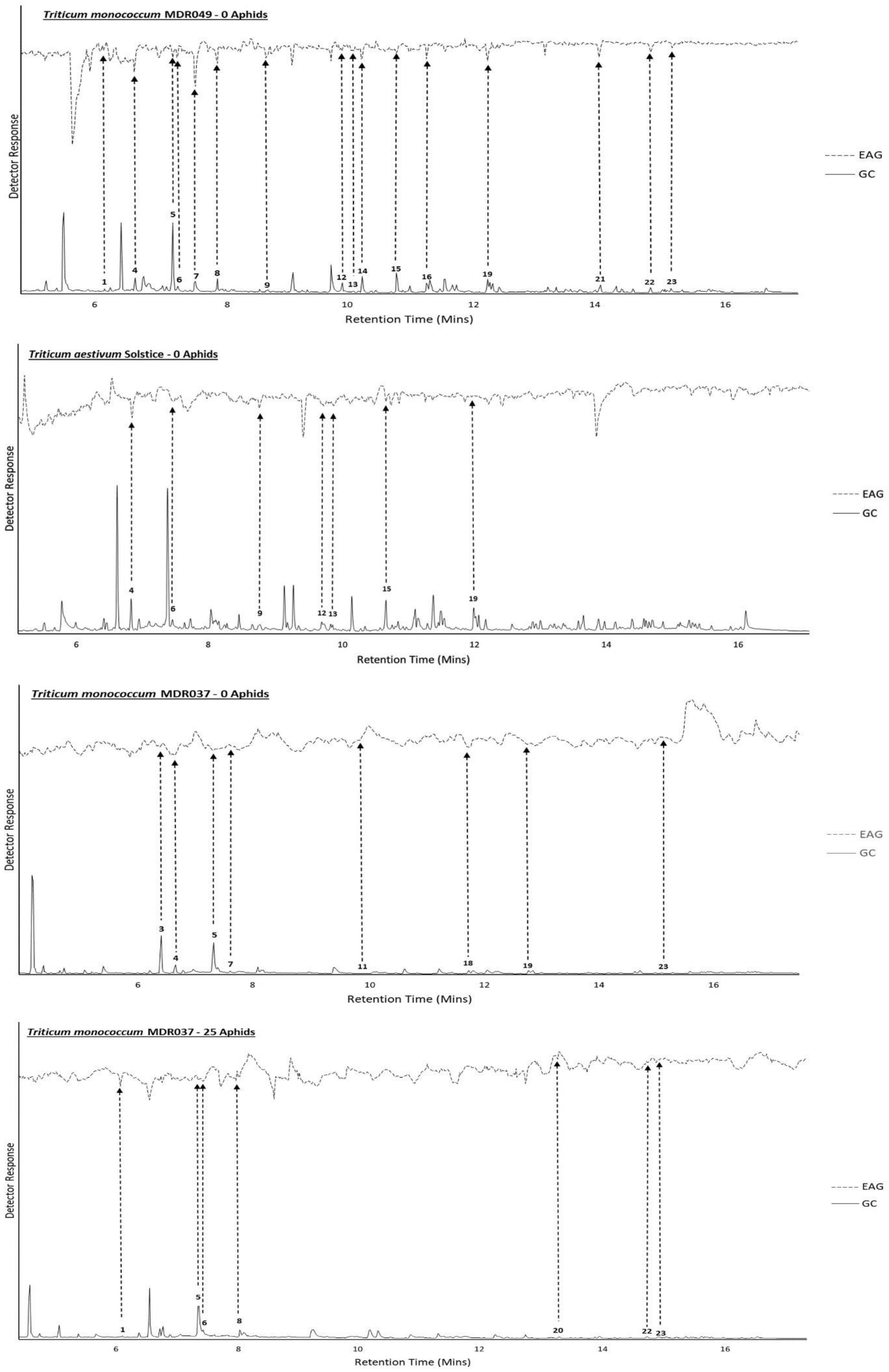

Triticum monococcum MDR049 - 0 Aphids
Triticum aestivum Solstice - 0 Aphids
Triticum monococcum MDR037 - 0 Aphids
Triticum monococcum MDR037 - 25 Aphids
Detector Response
Retention Time (Mins)
EAG
GC

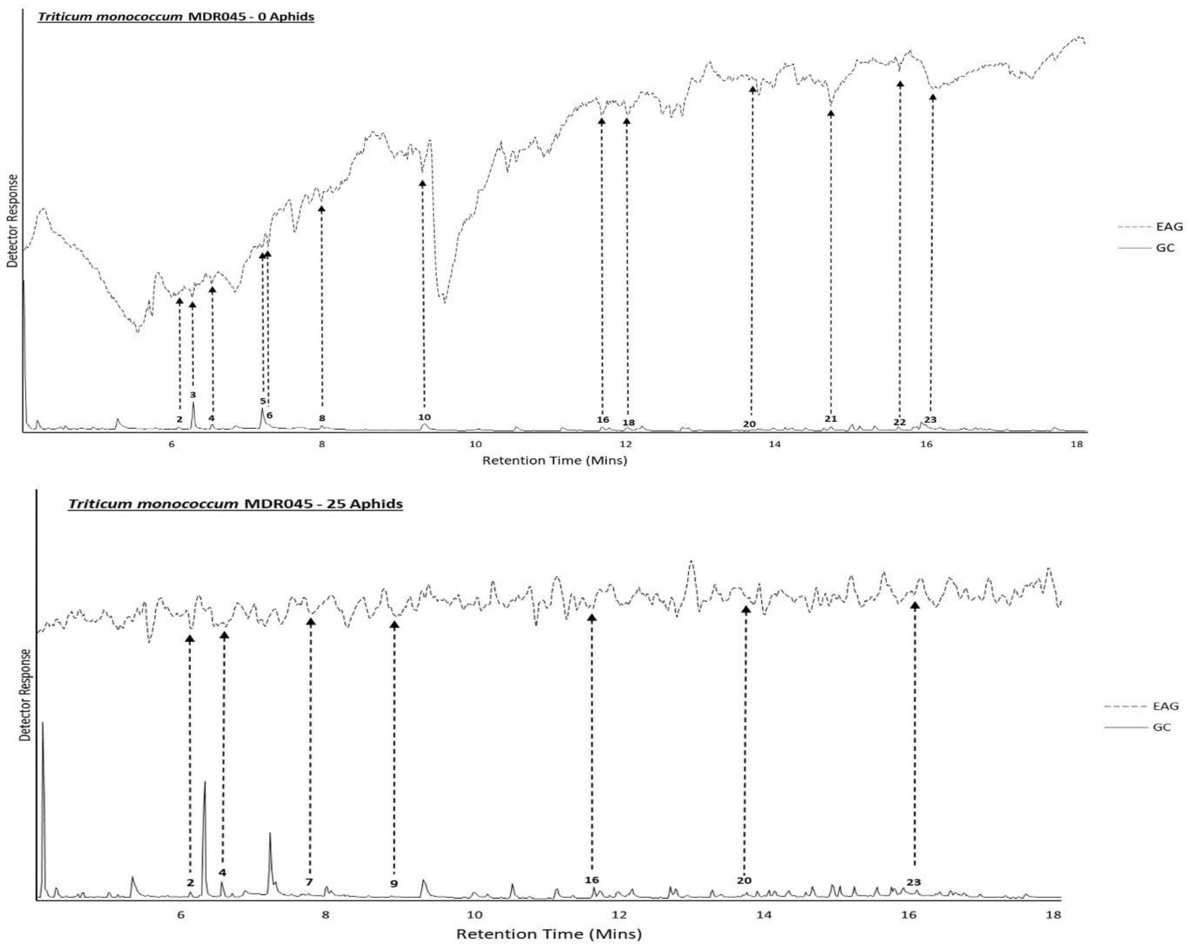


**Figure S2** Representative coupled GC-EAG traces showing antennal responses of alate grain aphids, *Sitobion avenae*, against headspace extracts collected from *S. avenae* (n=0)-infested *T. monococcum* MDR049 and *T. aestivum* Solstice and *S. avenae* (n=0, 25)-infested *T. monococcum* MDR037 and MDR045 for 24 hr. Upper trace, response of antenna; lower trace, FID response. GC peak numbers correspond to compounds listed in Table 1 with arrows indicating their respective EAG peak. Identifications confirmed by GC peak enhancement using authentic standards.

2

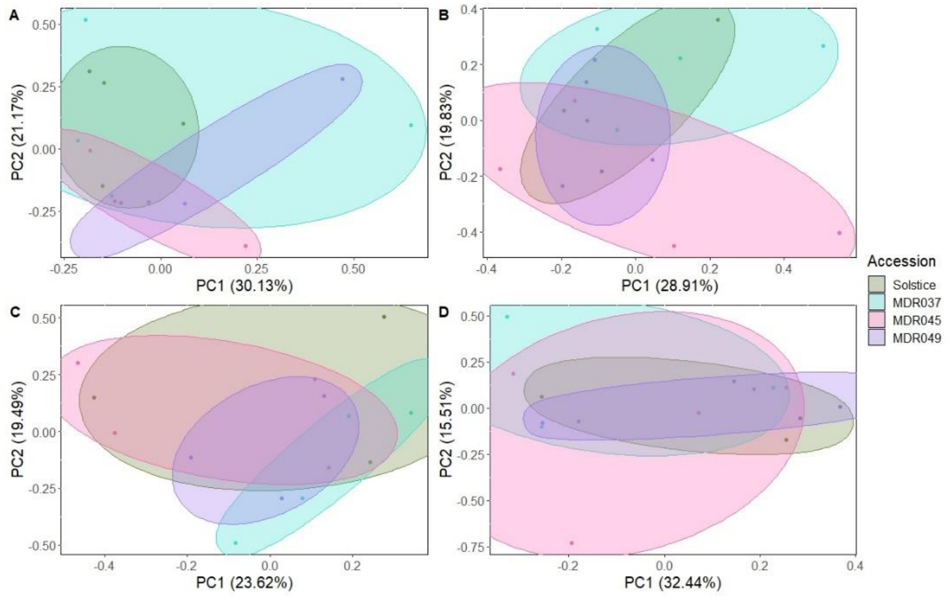


**Figure S3.** Principal component analysis (PCA) comparing EAG active VOC composition between *S. avenae* (n=0, 1, 5, 10, 25)-infested *T. monoccocum* MDR037, MDR045, MDR049 and *T. aestivum*. Ellipses represent the 95% confidence limits at each accession. Adonis permutation analysis provided P > 0.05 across all treatment combinations.
